# Taxonomic, geographic, and phylogenetic patterns in the conservation status of the squamate reptiles (Reptilia: Squamata) of Colombia

**DOI:** 10.1101/2025.11.18.689145

**Authors:** Juan D. Vásquez-Restrepo, Daniela García-Cobos

## Abstract

In this study, we evaluate for the first time the taxonomic, geographic, and phylogenetic patterns of both threatened and endemic squamate reptiles in Colombia at a broad scale. We employed community phylogenetic diversity metrics to assess patterns of species richness, endemism, and extinction risk. Additionally, we used spatially explicit autoregressive models to explore the relationships between this diversity and various abiotic and biotic variables. The data indicate that most reptile species have been evaluated by IUCN Red List specialist groups, although many assessments are outdated. Furthermore, there is a significant delay in local species assessments. We also observed that species richness is highest in the Amazonian region, followed by the Pacific-Andean transition. Interestingly, these richness distribution patterns do not necessarily align with the areas where most threatened and endemic species are found.

Specifically, our analysis identifies the Andean region as harboring the greatest number of endemic and threatened species, as well as exhibiting high turnover and phylogenetic endemism. Regarding phylogenetic patterns, we found that threatened species contribute significantly to evolutionary history. However, threatened categories do not exhibit a phylogenetic pattern beyond what would be expected by chance. Finally, we discuss the implications of assessing the threatened status of biodiversity across multiple levels beyond simply species number, and provide a national list of species prioritized for conservation from an evolutionary viewpoint.

## 1. Introduction

Biodiversity is one of the most remarkable features of life on Earth, showcasing immense variation, from unicellular to multicellular organisms, across different degrees of organic complexity. However, it is well-known that this diversity is not evenly distributed across the planet (Hawkins 2001; Mannion 2020). Tropical regions tend to harbor a larger number of species than higher latitudes, and this pattern is consistently observed across several groups of vertebrates and plants (Jenkins et al. 2013; Raz et al. 2023). Despite this phenomenon being identified more than 200 years ago (Humboldt 1807), there is no single or simple explanation for it. Instead, it is attributed to a set of hypotheses related to a combination of non-mutually exclusive factors, such as: evolutionary time, area, net productivity, spatial heterogeneity, climate stability, mutation rates, competition, among many others (Pianka 1966; Fine 2015; Zhang et al. 2022). On the other hand, as a multidimensional concept, biodiversity can be decomposed into distinct facets, including taxonomic, phylogenetic, and functional dimensions (Colwell 2009; Devictor et al. 2010; Jarzyna and Jetz 2016; Ochoa-Ochoa et al. 2020), each contributing differently to our understanding of the organization and structure of biological assemblages and ecosystems. Therefore, conserving tropical biodiversity requires protecting not just many species, but multiple biodiversity facets.

In the context of conservation, understanding how the different dimensions of biodiversity are distributed across space and evolutionary time is crucial for directing efforts and allocating resources. Traditionally, conservation has relied on species as the currency unit (Humphries et al. 1995; Cardillo 2023), prioritizing the number of species (taxonomic diversity) over their contributions to evolutionary history (phylogenetic diversity) or ecosystem maintenance (functional diversity). This approach likely persists because, in the absence of comprehensive data, taxonomic diversity serves as the “best” available surrogate, attempting to capture as much variation as possible. However, this “Noah’s Ark” strategy treats all species as equally valuable, ignoring redundancies and, consequently, the uniqueness of certain lineages. While phylogenetic and functional redundancies may enhance the resilience and stability of communities and ecosystems (Chave 2013; Ricotta et al. 2017; Biggs et al. 2023), including many species with the “same” ecological role or evolutionary history as priority for conservation could limit the ability of conservation interventions to capture the diversity of responses to environmental perturbations, thereby reducing the evolutionary potential of biological communities.

Tropical regions also exhibit the highest levels of sampling biases and gaps in taxonomic, phylogenetic, and geographic knowledge of species (Meyer et al. 2015; Hughes et al. 2021; Moura et al. 2024), resulting in a limited understanding of the impacts of anthropogenic disturbances and the vulnerability of species in these regions. These impacts, including habitat loss, climate change, pollution, and the introduction of invasive species, have intensified in recent decades, especially in tropical areas (Bellard et al. 2022). The magnitude and rapid occurrence of these disturbances within a short geological period have led to an acceleration of human-induced species losses in certain groups (Ceballos et al. 2015). Such human pressures, coupled with the rapid pace of global change, have contributed to dramatic biodiversity declines, often referred to as the “sixth mass extinction” (Ceballos et al. 2015; Barnosky et al. 2011). While many of these threats are well-documented, there is still a lack of comprehensive understanding regarding the vulnerability patterns of the affected species.

Due to its tropical location and complex geological history, Colombia is recognized as one of the most biodiverse countries in the world (Sarukhan and Dirzo 2013). Among this remarkable biodiversity, reptiles stand out as one of the most species-rich vertebrate groups (Páez et al. 2006). However, historical socio-economic inequities and political conflicts have hindered biodiversity research in certain regions while exacerbating specific threats (Clerici et al. 2020). For instance, albeit for the wrong reasons, the internal armed conflict of the past 60 years acted as a “shield” for natural ecosystems by limiting access to remote areas (Clerici et al. 2020; Prem et al. 2020), while simultaneously, making it difficult to carry out scientific research activities. Furthermore, Colombia has one of the highest rates of deforestation and illegal mining in the region (Armenteras et al. 2017; Schwartz et al. 2020; Wagner 2021). These activities are concentrated in areas with limited governmental presence, widespread poverty, insecurity, and significant social inequality (Etter et al. 2006; Gómez et al. 2021).

Although some studies have identified Colombia as one of the countries with the highest number of reptile species (Roll et al. 2016) and numerous threatened ones (Cox et al. 2022), a multi-faceted comprehensive local assessment is still lacking. Previous efforts to understand squamate distribution patterns at the national level include species checklists for snakes (Pérez-Santos and Moreno 1988) and lizards (Ayala and Castro, unpublished), as well as studies such as Sánchez et al. (1995) and Páez et al. (2006), which identified centers of endemism and species richness for approximately 490 squamate species based on regional bibliographic records. Nearly two decades after the last major assessment of national squamate distribution patterns, Colombia is now reported to host over 620 species of lizards and snakes (Uetz et al. 2024), highlighting the need for an updated and locally focused evaluation of this diverse group. Furthermore, no studies have yet evaluated spatial phylogenetic diversity to determine which regions of Colombia harbor distinct evolutionary lineages. This is a crucial gap, as such information could play a vital role in shaping local conservation priorities for squamates in this megadiverse country.

In this context, we aim to leverage all available information from various online sources to develop an updated geographical framework that improves our understanding of the conservation status of squamates in Colombia, thereby facilitating conservation initiatives and informing policy-making. Our specific objectives are to identify and describe geographical patterns of endemism and threats, and to analyze in the local context, for the first time, conservation status patterns within a phylogenetic framework.

## Methods

### 2.1. Taxonomic sampling

We compiled a list of squamate reptile species confirmed or potentially present in Colombia according to The Reptile Database (TRD: http://www.reptile-database.org) as of September 2023. Although this is a comprehensive list, it may include species erroneously attributed to Colombia or omit recent additions not yet incorporated, but serves as a practical proxy to explore general patterns (see Table S1). Moreover, since TRD is periodically updated, the total number of species used for the analyses may differ from the current figure. Our final dataset included 597 species of seven major monophyletic groups: Amphisbaenia (6 spp.), Anguimorpha (2 spp.), Gekkota (40 spp.), Gymnophthalmoidea (108 spp.), Iguania (110 spp.), Scincomorpha (7 spp.), and Serpentes (324 spp.). For each species, we gathered the following information: 1) IUCN Red List global assessment, based on the 2023-1 update (https://www.iucnredlist.org); 2) IUCN local assessment, as outlined in the most recent national red book (Morales-Betancourt et al. 2015); 3) national governmental conservation category, according to *Resolución 0126/2024* issued by the Colombian Ministry of Environment and Sustainable Development (MADS); 4) CITES appendix (https://cites.org); 5) endemism; 6) availability of geographic and phylogenetic data; 7) year of species description; and 8) year of the most recent IUCN Red List assessment.

### 2.2. Geographic sampling

We used the IUCN Red List distribution polygons filtered according to our species list, resulting in a total of 486 polygons, representing 81% of the species in our dataset. This total includes species in any category except for the non-evaluated ones. For spatial classification purposes, we adopted the Colombian natural regions division proposed by the *Instituto Geográfico Agustín Codazzi* (IGAC), which comprises five subregions: 1) Amazon (Amazon basin); 2) Andean (Andes and their inter-Andean valleys); 3) Caribbean (northern lowlands, Sierra Nevada de Santa Marta, and the archipelago of San Andrés, Providencia, and Santa Catalina); 4) Pacific (biogeographic Chocó); and 5) Orinoquia (Orinoco basin). To determine species endemism, we automatically identified species whose distributions did not extend beyond Colombia’s national boundaries (Table S2), allowing for a 5% error margin to account for commission errors (overprediction) in the polygons. Subsequently, we manually reviewed species without available distribution polygons to confirm endemism using distribution data from the literature.

### 2.3. Phylogenetic sampling

For analyses involving a phylogeny, we used the “fully-sampled” super-tree from Tonini et al. (2016). As their consensus tree is highly polytomous due to the mixed method used for its inference (DNA-based topology combined with taxonomic imputation of missing species), we subsampled 100 random trees to account for phylogenetic uncertainty (Rangel et al. 2015). These trees represent random dichotomizations of the consensus super-tree and were obtained from VertLife (https://vertlife.org). Previous studies have indicated that synthesis-based phylogenies provide robust results when calculating phylogenetic community metrics (Li et al. 2019). Therefore, all subsequent analyses were performed 100 times, using the average values derived from the frequency distribution of each parameter. In total, 551 species were included in the phylogenetic dataset, representing 92% of the species in our list.

### 2.4. Geographic analyses

Using species distribution polygons, we described both α and β diversities in geographical space. For α diversity, we generated a presence-absence matrix with the *lets.presab* function from the R package ‘letsR’ v.5.0 (Vilela and Villalobos 2015). For β diversity, we calculated mean community dissimilarity using the Sørensen index for each pixel, applying a 25-squared-pixels moving window. This was achieved with the *betaDiversity_speciesRaster* function from the ‘speciesRaster’ v.1.0 package (Title 2022). Additionally, we spatialized community phylogenetic metrics to summarize both phylogenetic endemism and overall evolutionary history (see section *2.5 Phylogenetic analyses*). Geographic analyses were conducted at a coarse grain (0.5° ≈ 55 km), but diversity metrics were also calculated at a finer resolution (0.16° ≈ 18.5 km), aiming to better understand their relationships with elevation by reducing variation in heterogeneous orographic regions like the Andes. Since phylogenetic endemism is sensitive to spatial resolution, we disaggregated the coarse-scale layer using the neighbor joining method instead of calculating it directly at a finer resolution. Coarse scales are generally more reliable for this metric due to the impact of commission errors in species distribution areas at finer grains (Daru et al. 2020a).

To explore the relationship between reptile diversities and environmental variables, we employed global spatial autoregressive models (see below), which account for the high spatial autocorrelation observed in species diversities (Table S3). The variables included in our analysis were as follows: elevation (Yamazaki et al. 2017); annual mean temperature [AMT], annual mean precipitation [AMP], isothermality [Iso], temperature seasonality [TSeaso], and precipitation seasonality [PSeaso] (Fick and Hijmans 2017); evapotranspiration [EvoT] (Zomer et al. 2022); forest cover (all vegetation taller than 5 m) [FC] (Hansen et al. 2013); and a topographic complexity index [TCI], calculated as the scores of a PCA incorporating elevation, terrain profile curvature, terrain tangential curvature, terrain roughness, and terrain slope (Amatulli et al. 2018). These variables were selected because they are most likely to influence the eco-physiology and diversity of squamates (Araújo et al. 2008; Powney et al. 2010; Qian 2010; Skeels et al. 2019). To address multicollinearity issues, we used a stepwise procedure to calculate the variance inflation factor (VIF) with the ‘usdm’ v.2.1.7 package (Naimi et al. 2014), applying the default threshold. As a result, elevation was excluded due to its high correlation with annual mean temperature.

We applied two types of spatial autoregressive models: the simultaneous autoregressive spatial lag (SAR_lag_) and the simultaneous autoregressive spatial error (SAR_error_). The SAR_lag_ model assumes that spatial dependence arises from the influence of neighboring locations, whereas the SAR_error_ model assumes that spatial dependence is due to unobserved variables (Dormann et al. 2007). The best-fitting model was selected based on the Akaike Information Criterion (AIC). Additionally, to evaluate whether the threatened categories (VU, EN, CR) were associated with endemicity, we performed a chi-squared (χ²) test and calculated the Cramer’s V coefficient. Cramer’s V measures the strength of association between two nominal variables, ranging from 0 (no association) to 1 (complete association).

### 2.5. Phylogenetic analyses

To investigate the degree of threat within an evolutionary context^1^, we followed the methodology outlined in Molina-Venegas et al. (2020) for analyzing phylogenetic patterns of extinction risk (see *op. cit.* for further details). First, we calculated phylogenetic diversity (PD_local_) across IUCN Red List categories using Faith’s PD (Faith 1992), and distance-relatedness metrics such as the Mean Nearest Taxon Distance (MNTD_local_) and the Mean Pairwise Phylogenetic Distance (MPD_local_). These calculations were performed using the *pd.query*, *mntd.query*, and *mpd.query* functions from the ‘PhyloMeasures’ v.2.1 package (Tsirogiannis and Sandel 2015). A major difference between MNTD and MPD lies in their sensitivity, while MNTD is more sensitive to patterns near the tree tips, making it suitable for analyzing extinction risk, MPD is better suited for describing overall phylogenetic diversity across spatial scales (Tucker et al. 2017). Moreover, since Faith’s PD is influenced by species number, we used it only to quantify the average contribution of species to each threat category.

Phylogenetic diversity was also z-standardized (indicated by the prefix ses-), based on null distribution generated by shuffling taxon labels within the phylogeny 999 times. This standardization accounts for variations in species richness among IUCN Red List categories and bioregions, improving interpretability. Values below -1.96 or above +1.96 are considered statistically significant (at α = 0.05). This approach allowed us to determine whether specific threat categories or regions exhibited a tendency for phylogenetic clustering (negative values) or dispersion (positive values). We also calculated the D statistic (Fritz and Purvis 2010) to assess the phylogenetic signal of binary traits (threatened vs. not threatened species), using the *phylo.d* function in the ‘caper’ v.1.0.3 package (Orme et al. 2023). A value near 0 indicates phylogenetic signal (non-random evolution), while values approaching 1 suggest a random evolutionary pattern.

Finally, we calculated phylogenetic endemism (Rosauer et al. 2009), which identifies regions where significant components of phylogenetic diversity are geographically restricted. This metric combines traditional measures of phylogenetic diversity with weighted endemism, providing insights into the spatial confinement of evolutionary lineages. For this last, we used the function *phylo_endemism* from the ‘phyloregion’ v.1.0.8 package (Daru et al. 2020b).

### 2.6 Species prioritization

To quantitatively identify species requiring prioritized conservation efforts, we employed the Evolutionarily Distinct and Globally Endangered (EDGE_local_) metric in its second version (Gumbs et al. 2023). The EDGE score integrates two key components: evolutionary distinctiveness (ED) and global endangerment (GE), collectively portraying a species’ “value and risk”. This metric provides a quantitative framework for prioritizing species that maximize future biodiversity through a cost-benefit approach (Isaac and Pearse 2018). The EDGE analysis prioritizes species that are both evolutionarily distinct (valuable from a phylogenetic perspective) and globally endangered (at high risk of extinction). Following the protocol outlined in Gumbs et al. (2023), we accounted for uncertainty in extinction risk data by using the default distribution of probabilities of extinction. Species were categorized into five groups based on their EDGE scores: 1) not EDGE (non-prioritized); 2) EDGE watch list (non-endangered species, but crucial for securing a large amount of phylogenetic diversity deep in the tree); 3) EDGE research list (species lacking sufficient data, prioritized for extinction risk research); 4) borderline EDGE species (species prone to prioritization, valuable in groups with high phylogenetic uncertainty); and 5) EDGE species (prioritized species due to high extinction risk and phylogenetic diversity).

## 3. Results

### 3.1 Taxonomic patterns

Up to the 2023-1 update, the IUCN Red List global assessment accounts for 92% of the squamate reptiles included in our dataset for Colombia, while the local categorization for 63% (excluding non-evaluated ones). These results include all species described up to 2023 for the global assessment and only those described up to 2013 for the local assessment (year in which the Colombian IUCN categorization workshop for reptiles took place). Most squamates are classified as LC, followed by DD and NE, with a smaller proportion categorized as NT or within a threat category (Fig. 1A–B). These last ones encompass 4.5% of the total, including those in VU, EN, and CR categories. This trend is consistent both globally and locally, but when considering endemic species only, the DD category slightly outnumbers LC species. Remarkably, endemic species account for 32% of the total squamate diversity in Colombia.

**Figure 1.**
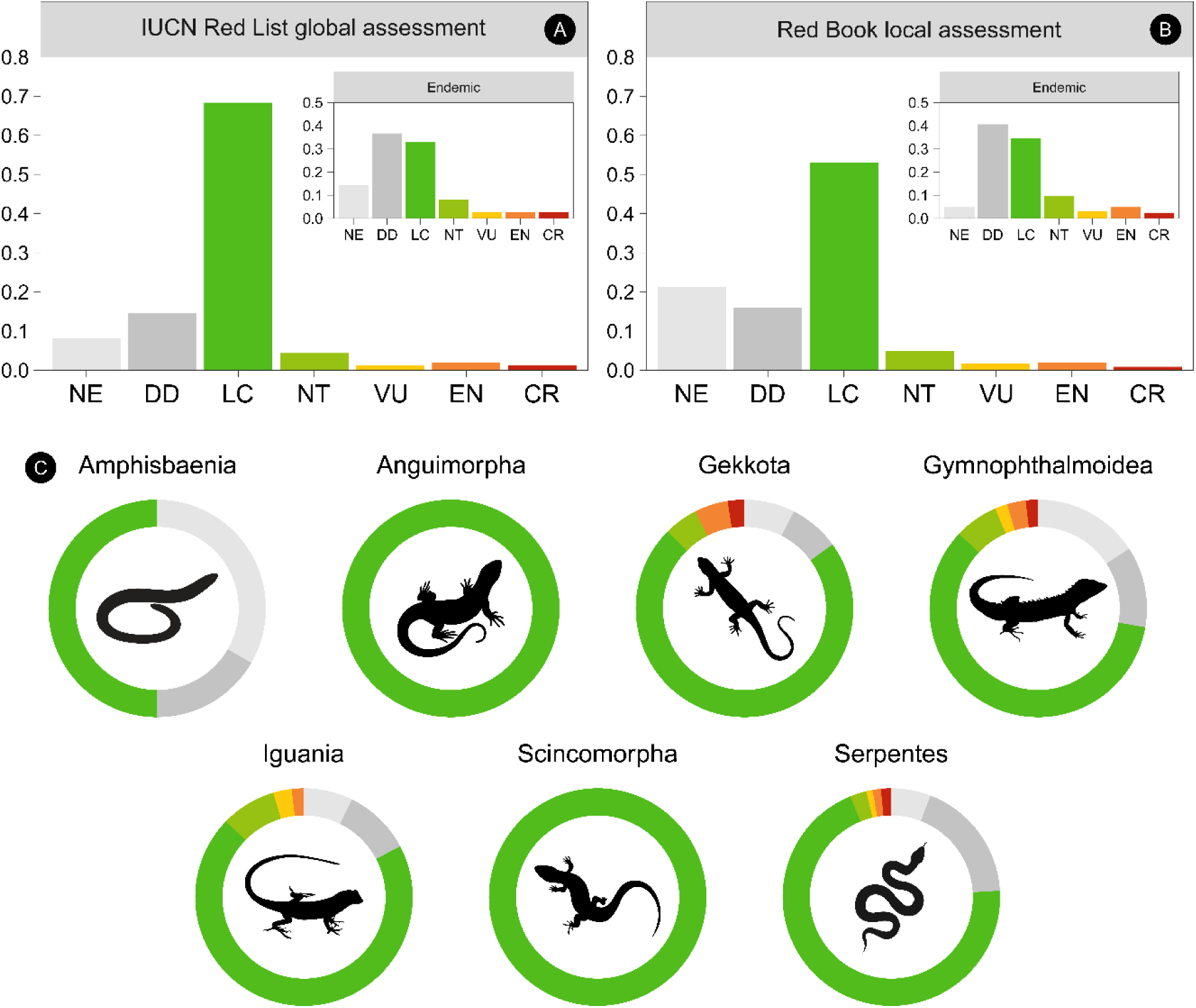
Summary of the conservation status of the Colombian squamate reptiles. **A:** global assessment. **B:** local assessment. **C:** taxonomic patterns following major monophyletic clades.

Within the seven major radiations in Squamata (Fig. 1C), 100% of the species in Anguimorpha and Scincomorpha are classified as LC. The only other group without any species in a threatened category is Amphisbaenia, with 50% classified as LC and the other half as NE or DD. For Gekkota, Gymnophthalmoidea, Iguania, and Serpentes, most species are considered LC, followed by DD and NE, with a small proportion (3.7% to 7.5%) in a threatened category. Among the 27 squamates globally classified as threatened, only one species (*Coniophanes andresensis*) is entirely insular.

Out of the 26 species classified as threatened (VU, EN, CR) in the local assessment, eight differ from the IUCN Red List global assessment: two shifted to DD, three moved out of a threatened category, two shifted to a higher-threat category, and one moved to a lower-threat category (Table 1). Additionally, six species classified as NE or DD in the local assessment were categorized as threatened in the global IUCN Red List assessment. Regarding CITES, 19 squamate reptiles from five families are listed under Appendix II (not necessarily threatened but subject to controlled trade). These include all boas and dwarf boas (Boidae and Tropidophiidae), certain snakes (Colubridae), and some iguanas and tegus (Iguanidae and Teiidae).

**Table 1.** Differences in extinction risk categories between the local (rows) and global (columns) IUCN Red List assessments. Values in the diagonal state instances where local and global assessment agree in the extinction risk category. Discrepancies between threatened species in the local assessment (red book) and the global IUCN Red List assessment are highlighted in bold.

|  | NE | DD | LC | NT | VU | EN | CR |
| --- | --- | --- | --- | --- | --- | --- | --- |
| NE | 45 | 14 | 80 | 4 | 0 | **3** | **1** |
| DD | 3 | 67 | 20 | 1 | 0 | **1** | **1** |
| LC | 1 | 1 | 301 | 0 | 0 | 0 | 0 |
| NT | 0 | 3 | 5 | 20 | 0 | 0 | 0 |
| VU | 0 | 0 | **2** | **1** | 7 | 0 | 0 |
| EN | 0 | **1** | 0 | 0 | **1** | 7 | **2** |
| CR | 0 | **1** | 0 | 0 | 0 | 0 | 4 |

**Table 2.**
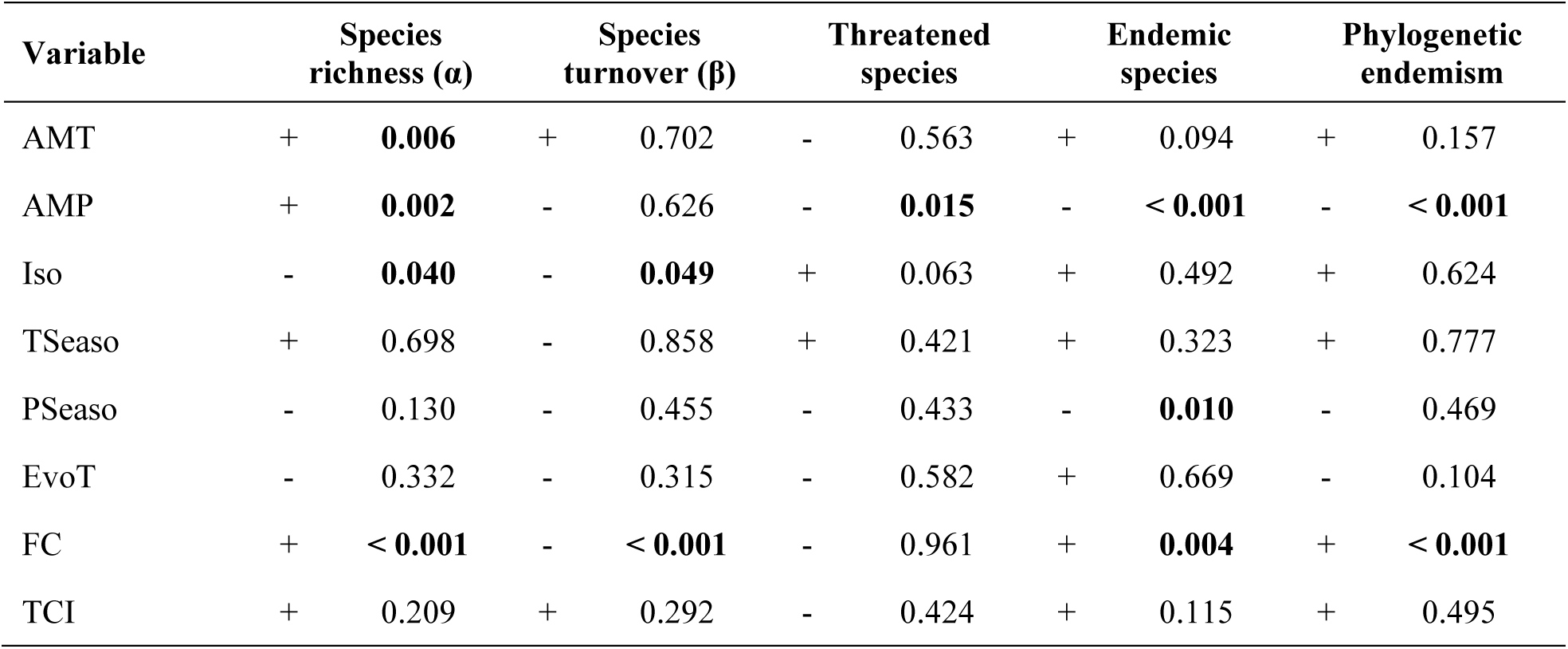
Effect of the environmental variables (as covariates) on the squamates diversity from the spatial autoregressive models (values correspond to *p-values* and significant ones are highlighted in bold). Variables are defined as: AMT (annual mean temperature), AMP (annual mean precipitation), ISO (isothermality), TSeaso (temperature seasonality), PSeaso (precipitation seasonality), EvoT (evotranspiration), FC (forest cover), TCI (topographic complexity index).

| Variable | | Species richness ($\alpha$) | | Species turnover ($\beta$) | | Threatened species | | Endemic species | | Phylogenetic endemism |
| --- | --- | --- | --- | --- | --- | --- | --- | --- | --- | --- |
| AMT | + | **0.006** | + | 0.702 | - | 0.563 | + | 0.094 | + | 0.157 |
| AMP | + | **0.002** | - | 0.626 | - | **0.015** | - | **< 0.001** | - | **< 0.001** |
| Iso | - | **0.040** | - | **0.049** | + | 0.063 | + | 0.492 | + | 0.624 |
| TSeaso | + | 0.698 | - | 0.858 | + | 0.421 | + | 0.323 | + | 0.777 |
| PSeaso | - | 0.130 | - | 0.455 | - | 0.433 | - | **0.010** | - | 0.469 |
| EvoT | - | 0.332 | - | 0.315 | - | 0.582 | + | 0.669 | - | 0.104 |
| FC | + | **< 0.001** | - | **< 0.001** | - | 0.961 | + | **0.004** | + | **< 0.001** |
| TCI | + | 0.209 | + | 0.292 | - | 0.424 | + | 0.115 | + | 0.495 |

### 3.2 Temporal patterns

The temporal pattern of species assessments indicates that most species described before the 2000s have been evaluated using the IUCN Red List criteria and categories (Fig. 2A). However, a significant deficit in species assessments is observed among those described within the last 15 years. Additionally, the proportion of species categorized as DD has consistently been higher for species described from the 1900s to the present. The average time between a species’ description and its IUCN Red List assessment is highly variable (20 ± 15 years), considering only species described after 1968, when the first reptile assessment was published (Honegger 1968). For squamates, the time between assessment and online publication can extend up to 10 years, though for most species this period is below six years (4.4 ± 2.1 years) (Table S1). Moreover, more than 50% of the squamate reptile species occurring in Colombia are flagged as “needs updating” (see Table S1), a tag that is applied to species whose assessments are 10 or more years old.

**Figure 2.**
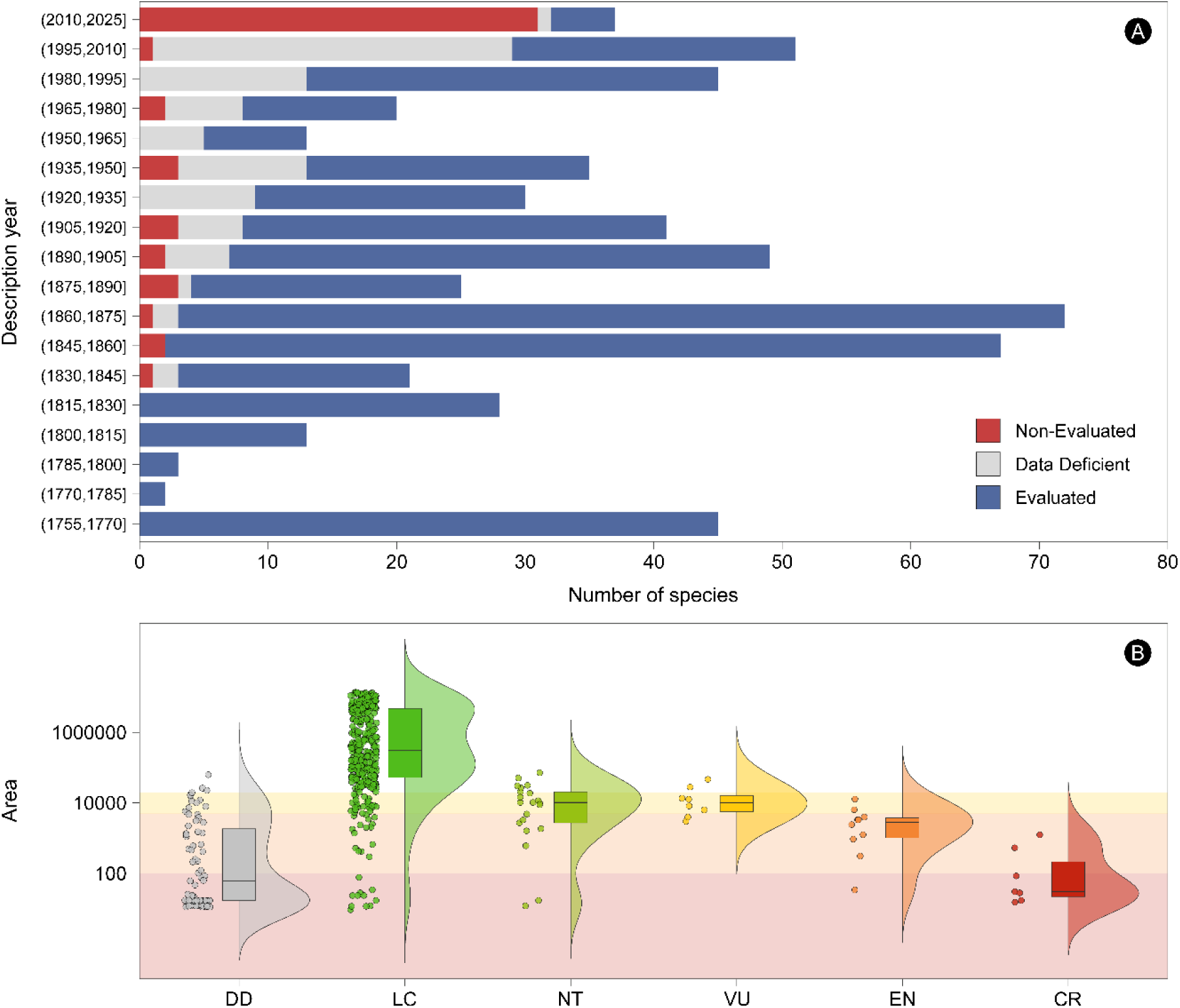
A: temporary pattern of extinction risk assessment regarding the species’ description year (in 15 years bins). **B:** species distribution areas (in km^2^) according to their IUCN Red List categories. The shadow areas indicate the Vulnerable (VU), Endangered (EN), and Critical Risk (CR) EOOs thresholds.

### 3.3 Geographic patterns

Most of the species in our dataset were assessed based on the IUCN Red List geographic criteria, primarily using their extent of occurrence (EOO), and to a lesser extent, their area of occupancy (AOO). However, we identified some species with EOOs that correspond to a different risk level or, in some cases, threatened species with EOOs outside the thresholds for their current category (Fig. 2B). Similarly, we observed species classified as LC with EOOs falling within the thresholds for threatened categories.

The bioregions with the highest species richness of squamate reptiles in Colombia are the Amazon, Orinoquia, Pacific, and the Andes, particularly in the Pacific-Andes transition zone (Fig. 3A). The highest species turnover, indicating the most dissimilar communities compared to their surroundings, is concentrated along the Pacific coast (Fig. 3B). However, this pattern is largely driven by the dominance of a single species, *Hydrophis platurus*, in Pacific marine communities. Excluding the marine communities, the regions with the highest species turnover shift to the Andes, Pacific, and the northernmost portion of the Caribbean, including the archipelago of San Andrés, Providencia, and Santa Catalina (Fig. 3B).

**Figure 3.**
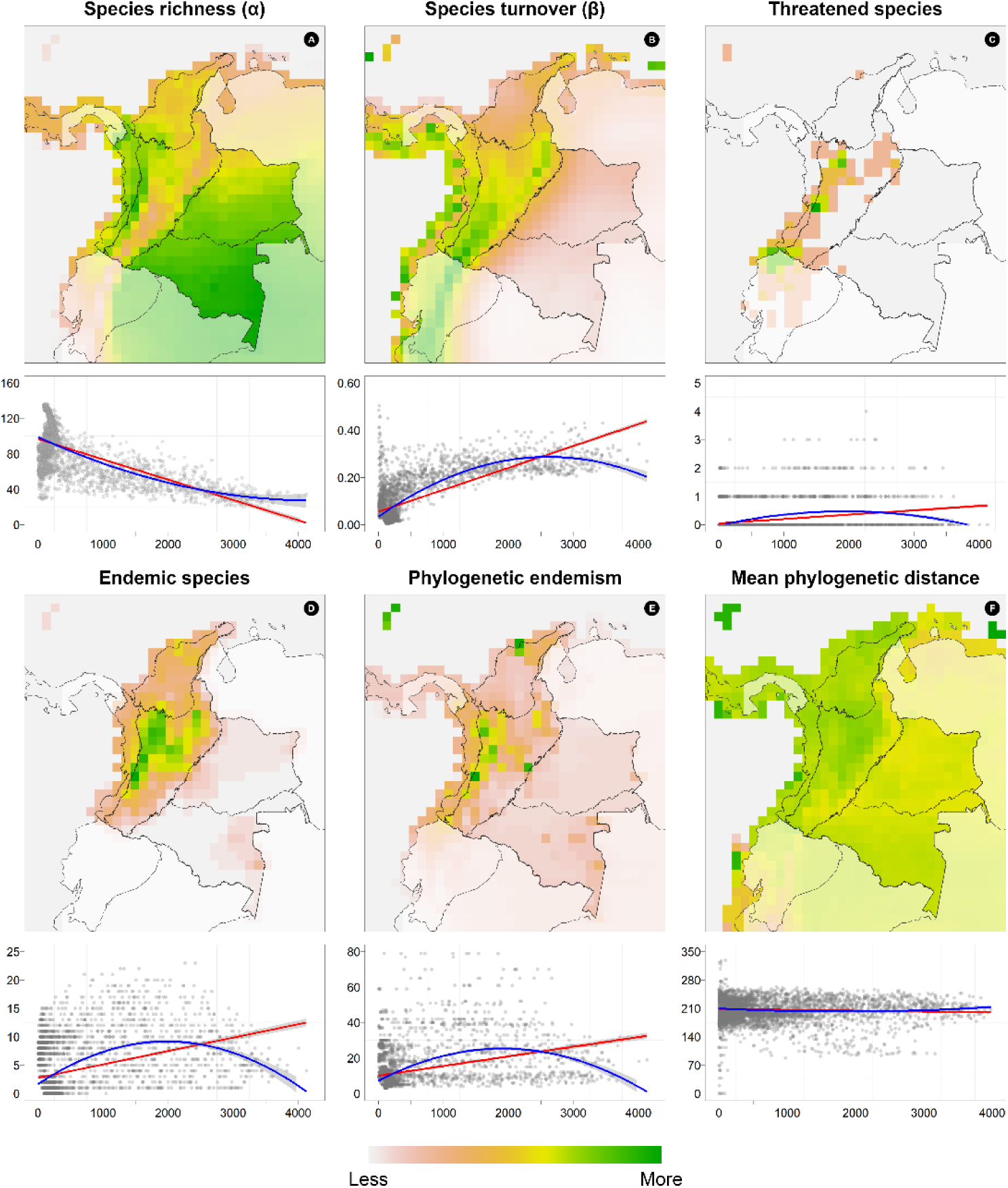
Geographic patterns of squamate reptiles’ diversity in Colombia. **A:** species richness. **B:** species turnover^2^. **C:** threatened species. **D:** endemic species. **E:** phylogenetic endemism. **F:** mean phylogenetic distance. Plots below maps show the relationship between the number of species and elevation (in meters). Trend lines in red correspond to linear models while those in blue to quadratic models (see Table S4 for performance metrics). Raster files are available as supplemental files.

The geographic distributions of threatened and endemic species show some overlap, with the majority found in the western and northern Andean region, the Pacific-Andes transition, and the Sierra Nevada de Santa Marta in the Caribbean (Fig. 3C–D). At a finer scale, threatened squamates are concentrated in three focal areas: 1) the northern portion of the Cordillera Central in the Antioquia department; 2) along the Cordillera Occidental and its adjacent lowlands in the central portion of Valle del Cauca department; and 3) the southernmost Colombian Andes in the department of Nariño.

Phylogenetic endemism also shows a distinct pattern, with higher values in the Andean and Pacific regions, the northernmost Caribbean, and specific hotspots like the Sierra Nevada de Santa Marta and the archipelago of San Andrés, Providencia, and Santa Catalina (Fig. 3E). In contrast, the average evolutionary contribution of species is slightly higher in the Andean, Caribbean, and Pacific regions compared to the Amazon and Orinoquia bioregions (Fig. 3F).

Almost all squamate diversity metrics showed an association with elevation (Fig. 3A–E) at different extents. For example, species richness exhibited a negative linear relationship with elevation, while most other metrics displayed a positive quadratic (hump-shaped) relationship (Table S4). Specifically, species richness decreased from lowlands to highlands, whereas species turnover tends to have higher values at intermediate elevations (approx. 1500–2500 m). Similarly, endemic species and phylogenetic endemism also exhibit a tendency toward higher values at mid elevations, though their variation remains considerably high. Despite being statistically significant, the rate of change and explained variance for both threatened species and MPD_local_ were negligible, indicating minimal response to elevation.

In terms of total species and endemicity proportions, the Andean region had the highest values, followed by the Pacific region, which had a high number of species but an intermediate proportion of endemics. The Caribbean region had an intermediate number of species but high endemism, while the Orinoquia and Amazonia regions had the lowest proportional values for both species richness and endemism (Fig. 4A). Species composition revealed two major groups (Fig. 4B), one consisting of the Amazonia and Orinoquia regions, and another comprising the Andes, Caribbean, and Pacific regions.

**Figure 4.**
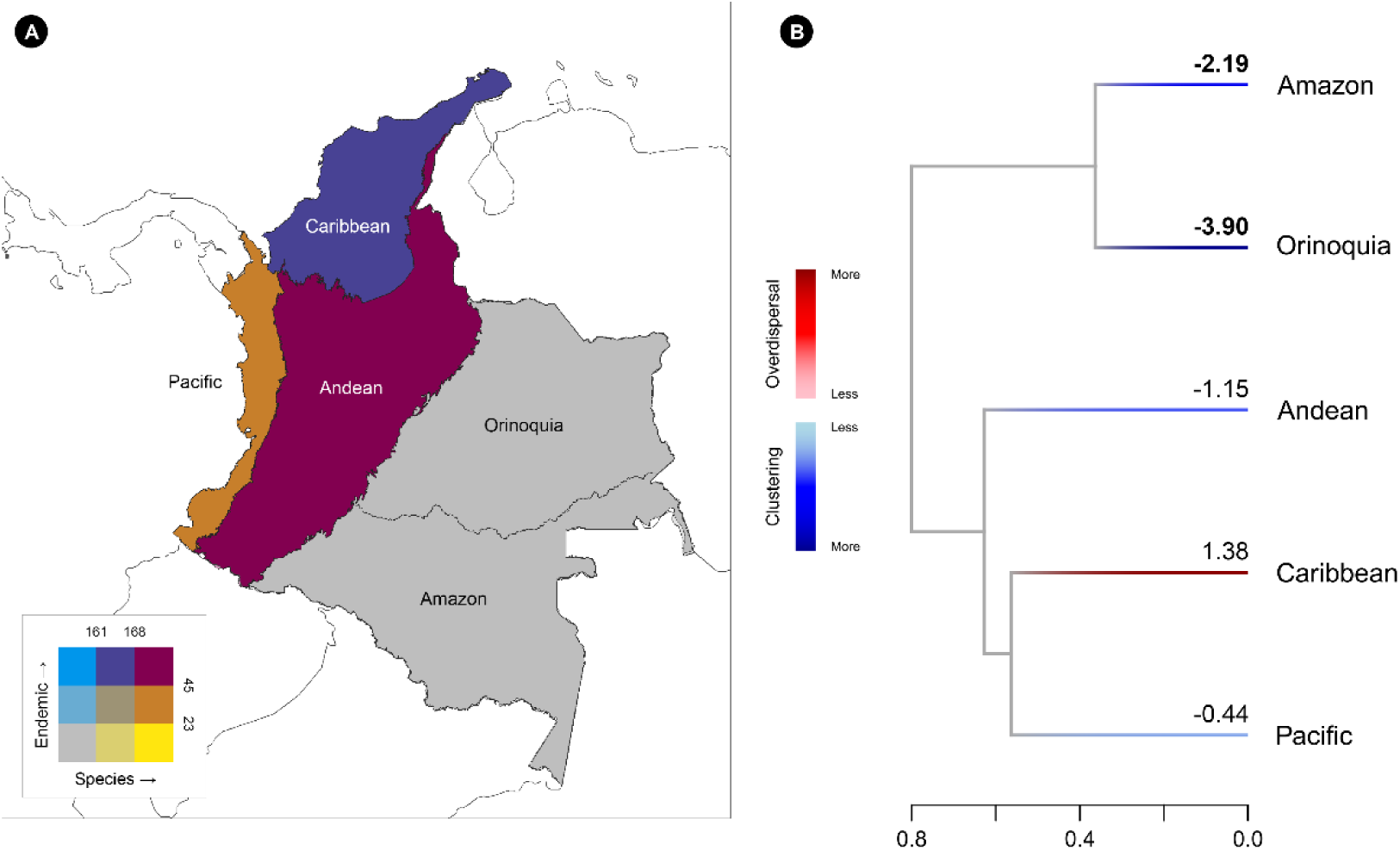
A: endemism per bioregion (quantile-based). **B:** bioregions Jaccard’s dissimilitude dendrogram. Branches are colored according to their sesMNTD_local_ values (numbers above branches). Significant patterns differing from chance (over or below ±1.96 threshold) are highlighted in bold.

### 3.4 Association with environmental variables

The χ² test revealed no significant relationship between threatened status and endemism (χ² = 0.22, df = 2, *p* = 0.89), with the Cramer’s V coefficient (CV = 0.07) further indicating a weak association between these variables. After accounting for spatial autocorrelation, the SAR_error_ model provided the best fit for most relationships, except for species richness, where the SAR_lag_ model performed better (Table S5). Species richness showed significant positive associations with temperature (AMT), precipitation (AMP), and forest cover (FC), and a significant negative relationship with isothermality (Iso). For species turnover, significant negative correlations were found with both isothermality and forest cover. Threatened species exhibited a significant negative association with precipitation, while endemic species showed negative correlations with both precipitation and precipitation seasonality (PSeaso), along with a positive association with forest cover. For phylogenetic endemism, precipitation had a significant inverse association, while forest cover showed a positive correlation.

### 3.5 Phylogenetic patterns

Among the five bioregions, only the Amazon and Orinoquia exhibited phylogenetic patterns significantly different from chance, both showing a trend toward phylogenetic clustering (Fig. 4B). For squamate reptiles in Colombia, the average phylogenetic contribution per IUCN Red List category revealed that threatened species (VU, EN, CR) and near threatened species (NT) contribute significantly more to overall phylogenetic diversity than LC species, although their contribution is exceeded by NE species (Fig. 5A). Regarding the sesMNTD_local_ values, all IUCN categories except DD fell within the non-significant area (Fig. 5B), indicating that their phylogenetic distribution does not deviate from a random pattern (Fig. 6). This is further supported by the high D statistic value (D = 0.82 ± 0.08), suggesting that extinction risk as the manifestation of extrinsic and intrinsic populational features evolves as a random trait.

**Figure 5.**
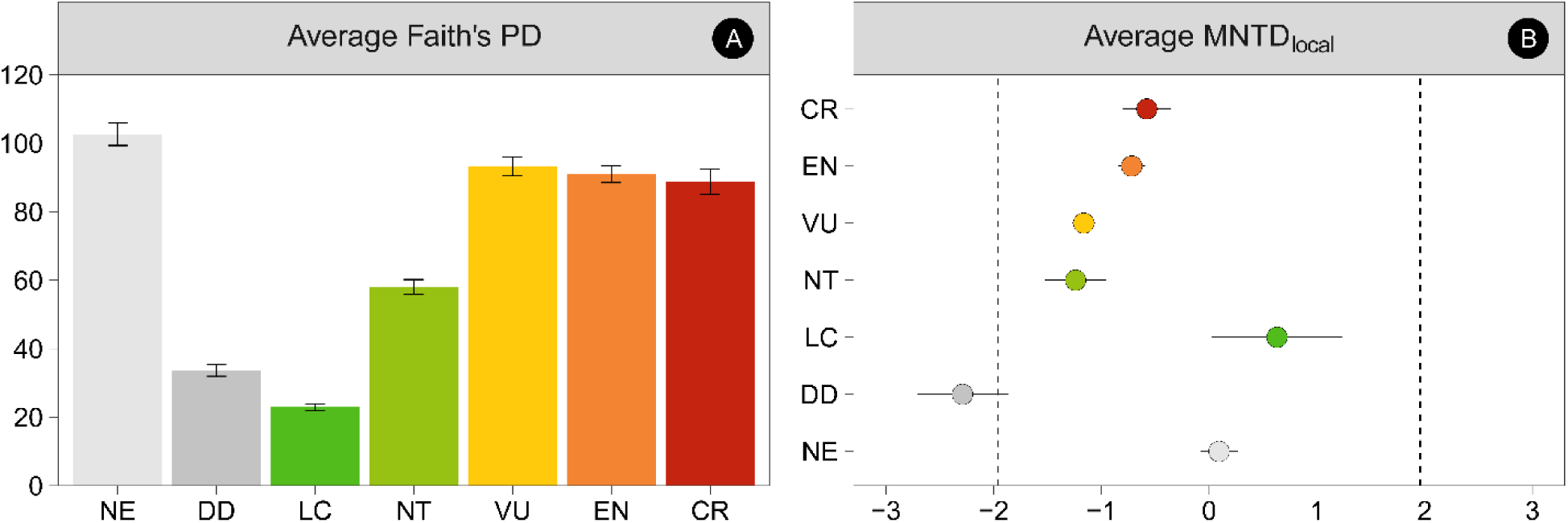
A: Average contribution of each IUCN Red List category to the PD_local_ (Faith’s PD). **B:** Standardized values (z-scores) for the MNTD_local_. Vertical dashed lines indicate the ±1.96 threshold of significance (⍺ = 0.05).

**Figure 6.**
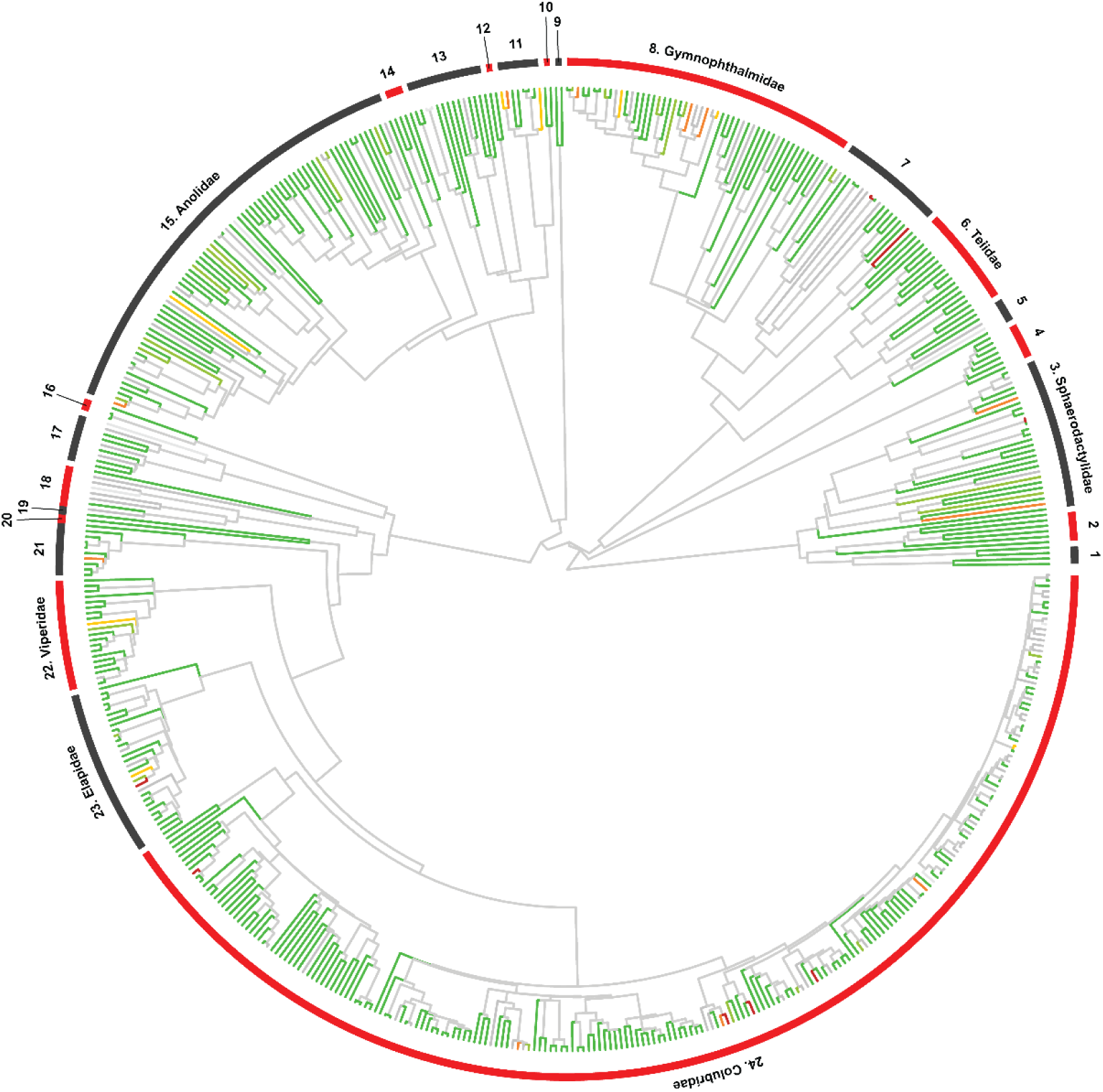
Pruned tree for the squamate reptile’s species occurring in Colombia. Branches are colored according to their conservation status following the IUCN Red List color codes. For illustration purposes, we depicted a random tree from our 100 subsamples. 1: Phyllodactylidae, 2: Gekkonidae, 3: Sphaerodactylidae, 4: Scincidae, 5: Amphisbaenidae, 6: Teiidae, 7: Alopoglossidae, 8: Gymnophthalmidae, 9: Diploglossidae, 10: Polychrotidae, 11: Hoplocercidae, 12: Iguanidae, 13: Tropiduridae, 14: Corytophanidae, 15: Anolidae, 16: Typhlopidae, 17: Leptotyphlopidae, 18: Anomalepididae, 19: Aniliidae, 20: Tropidophiidae, 21: Boidae, 22: Viperidae, 23: Elapidae, 24: Colubridae

### 3.6 Species priority for conservation

The EDGE_local_ analysis identified 19 EDGE species and six borderline EDGE species (Table 3), along with 11 species on the EDGE research list, 11 on the EDGE watch list, and 504 species not classified as EDGE (Table S6). The prioritized squamate reptiles belong to 12 genera distributed across nine families: Alopoglossidae, Anolidae, Boidae, Colubridae, Elapidae, Gymnophthalmidae, Hoplocercidae, Sphaerodactylidae, and Viperidae. The number of species per EDGE genus is fairly equitable, with most genera containing 1–2 species. Since not all threatened species are EDGE species, the EDGE species collectively account for 90% (120.17 Myr) of the expected phylogenetic diversity loss of all threatened species (133.45 Myr).

**Table 3.** First 25 prioritizing species for conservation based on their EDGE_local_ scores (a complete list can be found in Table S6).

| Major clade | Family | Species | Median EDGE | IUCN |
| --- | --- | --- | --- | --- |
| Gekkota | Sphaerodactylidae | *Sphaerodactylus scapularis* | 15.142 | EN |
| Gekkota | Sphaerodactylidae | *Lepidoblepharis miyatai* | 12.347 | CR |
| Serpentes | Colubridae | *Emmochliophis miops* | 10.570 | CR |
| Serpentes | Colubridae | *Coniophanes andresensis* | 9.232 | CR |
| Iguania | Anolidae | *Anolis calimae* | 8.068 | VU |
| Gymnophthalmoidea | Alopoglossidae | *Alopoglossus danieli* | 6.786 | CR |
| Iguania | Anolidae | *Anolis ruizii* | 6.702 | EN |
| Gekkota | Sphaerodactylidae | *Lepidoblepharis williamsi* | 6.614 | EN |
| Gymnophthalmoidea | Alopoglossidae | *Alopoglossus lehmanni* | 6.587 | CR |
| Serpentes | Colubridae | *Synophis plectovertebralis* | 5.114 | CR |
| Serpentes | Elapidae | *Micrurus medemi* | 5.059 | CR |
| Iguania | Hoplocercidae | *Enyalioides groi* | 5.008 | EN |
| Iguania | Hoplocercidae | *Enyalioides oshaughnessyi* | 4.050 | VU |
| Serpentes | Boidae | *Corallus blombergi* | 3.792 | EN |
| Gymnophthalmoidea | Gymnophthalmidae | *Riama columbiana* | 3.217 | EN |
| Serpentes | Viperidae | *Bothrocophias campbelli* | 3.165 | VU |
| Serpentes | Colubridae | *Synophis bicolor* | 3.118 | EN |
| Iguania | Hoplocercidae | *Enyalioides annularis* | 3.051 | VU |
| Gymnophthalmoidea | Gymnophthalmidae | *Riama simotera* | 2.552 | EN |
| Serpentes | Colubridae | *Dendrophidion boshelli*<sup>1</sup> | 3.210 | CR |
| Serpentes | Colubridae | *Dipsas ellipsifera*<sup>1</sup> | 2.356 | EN |
| Gymnophthalmoidea | Gymnophthalmidae | *Anadia pamplonensis*<sup>1</sup> | 1.910 | EN |
| Gymnophthalmoidea | Gymnophthalmidae | *Andinosaura laevis*<sup>1</sup> | 1.426 | VU |
| Serpentes | Colubridae | *Saphenophis sneiderni*<sup>1</sup> | 1.392 | EN |
| Serpentes | Elapidae | *Micrurus sangilensis*<sup>1</sup> | 1.160 | VU |

<sup>1</sup>Borderline EDGE species.

Furthermore, safeguarding the top 100 highest-ranked species, regardless of their classification, would secure almost two-thirds of the expected phylogenetic diversity loss across all Colombian squamates (Table S6).

## 4. Discussion

### 4.1 Overview of the Colombian squamates threat status

Despite their remarkable diversity, the proportion of threatened squamates in Colombia remains relatively low, compared to other reptiles such as turtles, where over 42% of species are threatened (Páez et al. 2022). This disparity can be partly attributed to the greater historical pressures faced by groups like turtles and crocodiles, including extensive use, exploitation, and illegal trade (Rueda-Almonacid et al. 2007). However, a major limiting factor in accurately assessing extinction risk in squamates is their low detectability, secretive behavior, and low abundances in certain groups (Böhm et al. 2013). Only 23% of the squamate reptiles in Colombia are categorized as NE or DD, which is comparable to their global status. However, this is particularly concerning for endemic species, of which nearly 50% are listed in one of these two categories. On the other hand, 89% of the evaluated species were assessed over 10 years ago, which is officially considered outdated according to IUCN standards.

The geographic range, is one of the most important criteria used by IUCN Red List specialists to assess threat status. This reliance stems from the lack of data on other criteria, particularly those related to population dynamics (Le Breton et al. 2019). For example, ground snakes of the genus *Atractus*, comprising ca. 65 species in Colombia, lack sufficient data for about 50% of their species. Consequently, many species with restricted ranges or poorly known distributions, particularly among endemics, may remain overlooked in conservation prioritization despite their potential vulnerability. One way to address these data gaps is by encouraging a broader use of curated information during taxonomic work. Although taxonomic research is frequent, it often remains limited to the description or revision of taxa. The assessment workflow could be significantly optimized if taxonomists went beyond traditional practices and systematically made curated datasets available as part of the process (Tapley et al. 2018). A recent example illustrating this approach is the description of a new species of *Echinosaura* from Colombia and Panama, where the authors not only described the new taxon but also used the curated data to reassess the conservation status of all species within the genus (Vásquez-Restrepo and Daza 2025). In addition to these challenges in evaluating individual species, squamates continue to face significant threats, including habitat loss, climate change, and the introduction of invasive alien species (Todd et al. 2010). Therefore, it is crucial to address conservation challenges from multiple fronts.

As we showed, some species have distribution ranges that do not align with their current threat categories. While this does not mean that all these species should be immediately moved to another category, it serves as a good starting point for future evaluations (Table S7). The fact that the EOO of certain species does not match what would be expected based on their current threat category is not, by itself, definitive proof that a reassessment is necessary. When applying the IUCN Red List geographic criteria, it is important to consider not only the distribution area but also the number of known localities and at least one additional factor related to potential threats. If all required criteria are not met, the species does not qualify for listing under threatened category, even if two out of the three geographic subcriteria are fulfilled. For example, this is the case for two of the three lizard species endemic to Malpelo Island (*Anolis agassizi* and *Diploglossus millepunctatus*), which have highly restricted distributions and are known from a single locality, but whose populations are estimated to range from thousands to hundreds of thousands of individuals (López-Victoria et al. 2011).

### 4.2 Patterns of geographic diversity and endemism

Broad geographic patterns of reptile diversity in Colombia had not been revisited and updated since the 1990s (Sanchez et al. 1995), despite significant contributions from many researchers in updating species list on specific regions across the country (e.g., Carvajal-Cogollo and Urbina-Cardona 2008; Castro-Herrera and Vargas-Salinas 2008; Llano-Mejía et al. 2010; Pedroza-Banda et al. 2014; Restrepo et al. 2017; among others). Our findings support earlier studies indicating that the highest reptile species richness is concentrated in the Amazonian region (Costello et al. 2022) and the Pacific (Lynch et al. 1997). On the other hand, the Andean region in Colombia harbors the highest number of endemic squamates, consistent with previous studies highlighting this area as a hotspot of endemism (Duellman 1999; Kattan et al. 2004; Rahbek et al. 2019). Moreover, diversity in the Caribbean is highly influenced by the Sierra Nevada de Santa Marta, identified as one of the richest regions per unit area in Colombia, mirroring similar trends in other vertebrate and invertebrate groups (Kattan et al. 2004). This pattern can be explained, in part, by the higher levels of species turnover in mountain areas compared to lowlands, which also has been observed in other groups like birds, mammals, and amphibians (Tenorio et al. 2023). Turnover likely reflects the topographic and climatic heterogeneity in the Andes, restricting organismal dispersal (Kattan et al. 2004; Graham et al. 2014), providing ecological opportunities by novel high-elevation habitats (Graham 2009; Hoorn et al. 2010), or differentially promoting speciation and extinction rates (Graham et al. 2014).

Although we observed interesting geographic patterns, we acknowledge that species distribution ranges are approximations (Hortal et al. 2015), which may vary if a lineage hides cryptic diversity (Funk et al. 2011; Böhm et al. 2013) or due to differences in species delimitation concepts. Such factors can lead to more restricted distributions as taxonomists split or describe new species, influencing extinction risk assessments (Robuchon et al. 2019). Endemism patterns may also reflect biases in geographic and taxonomic sampling efforts. For instance, there has been a historical trend of describing more Andean frogs of the genus *Pristimantis* compared to other groups or regions (Reyes-Puig and Mancero 2022). Additionally, the Andean region not only concentrates a high proportion of Colombia’s population but also hosts major research centers, making it more accessible compared to isolated regions like the Amazon and Orinoquia (Camacho-Rozo and Urbina-Cardona 2024).

### 4.3 Environmental relationships

Overall squamate diversity appears to be favored by temperature (AMT). This supports the altitudinal richness gradient observed from lowlands to highlands, as expected based on the thermal physiology of reptiles in humid climates such as the Andes (McCain 2010). Under this scenario, thermophilization in mountain areas could exacerbate negative pressures on threatened and endemic species in the Andean region and the Sierra Nevada de Santa Marta.

Precipitation (AMP) also promotes overall species diversity but has the opposite effect on the richness of threatened and endemic species. This aligns with historical trends of higher pluviosity in the Pacific region, Amazon, and Orinoquia (where greater species richness is found) and lower precipitation rates in the Andes (where most endemic and threatened squamates are located) (Poveda et al. 2007). However, at least for species diversity, this pattern challenges the general trend observed in squamates, where higher diversity is typically expected in drier areas (McCain 2010; Pie et al. 2017). In mountain regions, it has been suggested that both temperature and precipitation may act together in shaping reptile diversity (McCain 2010). For threatened and endemic species, their distribution suggests a “low plateau with a mid-elevation peak” pattern, as expected in wet mountain environments (McCain 2010).

Additionally, endemism benefits from less seasonality in precipitation (PSeaso). This may relate to local adaptations that can persist over long periods when the environment remains stable. For instance, previous studies have found that more stable climates favor *in situ* diversification processes in groups such as amphibians (Ochoa-Ochoa et al. 2019). Conversely, greater seasonal variations in temperature (Iso) benefit both α and β diversities, possibly because they promote temporal niche separation.

As mentioned, some squamate reptiles may be challenging to assess and monitor, requiring the implementation of indirect conservation strategies focused on ecosystemic scales in the short term. Given the positive association found between overall squamate diversity and endemism (at both taxonomic and phylogenetic levels) with the presence of forest cover, our results support such indirect conservation strategies. However, it is also important to recognize that landscape heterogeneity promotes species turnover and, therefore, more diverse communities. Further studies are necessary to fine-tune these details.

### 4.4 Evaluating the extinction risk in a phylogenetic context

Categories reflecting extinction risk are not intrinsic traits of a species but rather, to some extent, reflect certain population attributes (Chichorro et al. 2022). Attributes such as birth rates, fecundity, abundance, dispersal capacity, and tolerance to disturbances influence geographical range size and population numbers, key aspects used in extinction risk assessments based on IUCN Red List criteria (Purvis et al. 2000; Chichorro et al. 2022). However, although, some populations of a particular species may have demographic parameters that are favorable for a low risk, extrinsic factors stemming from human intervention can alter its vulnerability. Analyzing extinction risk within an evolutionary context requires acknowledging that this is not a trait from which ancestry can be reconstructed. Patterns may be obscured by extrinsic factors unrelated to the phylogenetic history of lineages. Instead, it should be seen as a tool whose utility lies in understanding the role of the evolutionary contribution of lineages for our conservation actions.

### 4.5 Conservation from an evolutionary perspective

When diversity is understood as a multi-faceted variable (Burbano-Girón et al. 2022) a fundamental set of questions arise: what should we prioritize? Taxonomic diversity? Phylogenetic diversity? Functional diversity? The answer is far from simple, as it depends on various factors, ranging from research interests to methodological and budgetary constraints. This complexity is further exacerbated by the fact that these three facets of biodiversity often do not overlap geographically (Vásquez-Restrepo et al. 2022). Therefore, it is crucial to deconstruct diversity patterns to identify where conservation efforts should be concentrated to maximize their impact. The second question emerges when considering the trade-offs of preserving threatened species (Isaac and Pearse 2018): should we invest in a few species prone to extinction or in many species to prevent them from becoming prone to extinction? Beyond personal, emotional, or aesthetic considerations, the next question is: which species contribute the most to biodiversity, and how can efforts and resources be prioritized accordingly?

In the case of Colombian squamates, phylogenetic diversity provides a partial answer. Our analyses suggest that the average phylogenetic contribution of both threatened species (categorized as VU, EN, or CR) and Near Threatened species (NT) is greater than that of species categorized as Least Concern (LC). This indicates that preserving the relatively few threatened species could significantly enhance the overall phylogenetic diversity of squamate assemblages. However, the high contribution of Not Evaluated (NE) species to phylogenetic diversity suggests that this framework may need adjustment as more species are assessed. According to our results, the phylogenetic composition of squamate assemblages in the Andean, Caribbean, and Pacific regions is indistinguishable from random assemblages, masking hypotheses about the eco-evolutionary processes shaping their diversity. Nonetheless, this apparent randomness, combined with the significant contribution of species from these regions to the overall evolutionary history, highlights the uniqueness of these assemblages, which may be at risk from current and future threats.

As previously mentioned, any phylogenetic metric reliant on branch lengths is sensitive to tree topology, which can vary naturally (e.g., missing species) or artificially (e.g., tree pruning), often leading to a “push to the past” effect (Calderón del Cid et al. 2024; Vásquez-Restrepo and Diago-Toro 2024). This phenomenon tends to inflate branch lengths, causing them to no longer accurately reflect true lineage ages and leads to an overestimation of evolutionary contributions. While such distortions may cast doubt on the utility of evolutionary metrics for conservation unless applied at global scales (both geographic and taxonomic), the other side of the coin is that when all species are considered, local conservation priorities can be minimized, creating a false sense of security, as global and local priorities do not always align. For instance, while species like *Anolis ruizii* and *Lepidoblepharis miyatai* appear to be shared priorities between the global assessment (Gumbs et al. 2024) and our national evaluation, others such as *Anolis calimae*, *Coniophanes andresensis*, *Micrurus medemi*, and *Riama columbiana* are not (all of them endemics). Therefore, recognizing the scale-dependency of evolutionary metrics applied to conservation and interpreting them as local proxies can provide valuable insights at finer resolutions.

### 4.6 Threats to the Colombian squamates

The Andes and the Amazon, despite their extraordinary biodiversity, face anthropogenic pressures that threaten both their ecosystems and species. Accelerated forest loss and pollution from extractive activities such as illegal mining, fires, and timber logging are among the most pervasive ones (Armenteras et al. 2017; Schwartz et al. 2020; Wagner 2021). For instance, the Andean region, which harbors the highest numbers of endemic and threatened reptile species, is particularly vulnerable. This vulnerability is related with the high human population density, as approximately 80% of Colombia’s population (over 40 million people) resides in the Andes, which constitutes just 25% of the country’s total area (Ospina 2006; World Bank Group 2021). This intense human presence has transformed the region significantly, with around 60% of its natural cover lost since pre-Columbian times (Armenteras et al. 2011). Additionally, the tropical Andes are highly sensitive to climate change, with rising temperatures driving ecosystems to migrate upward and altering rainfall regimes in less predictable ways (Anderson et al. 2011; Báez et al. 2016).

### 4.7 Implications for conservation actions

The high proportion of endemic species classified as NE or DD poses a significant risk of overlooking unique elements specific to the country in conservation decision-making, thereby increasing the probability of their potential loss. Similarly, discrepancies between the ranges of some species and their current threat categories are a cause for concern. Furthermore, the identification of EDGE and borderline species offers valuable insights that can be effectively integrated into prioritization processes, such as those concerning protected areas. For example, similar to findings in a previous study (Moya-Bedoya et al. 2024), species like *Micrurus medemi* and *Bothrocophias campbelli* were identified as high-priority candidates for the conservation of venomous snakes in Colombia. Conservation strategies, such as those implemented by Key Biodiversity Areas (KBAs), could greatly benefit from using such tools to prioritize regions where biodiversity management efforts are most urgently needed. This approach is particularly relevant for methodologies like KBAs, which focus on conserving individual species rather than broader aspects such as landscapes or ecosystem functions.

## Data availability statement

Supplementary files included in this manuscript can be found on the publishers’ website.

## Acknowledgments

We would like to thank Elkin A. Tenorio, Nicolás Urbina, and Sandra P. Galeano for their valuable comments, which helped to clarify and improve several sections of this manuscript. We also thank the anonymous reviewers for their feedback, which enhanced the manuscript’s readability and clarity.

## Footnotes

^1^All phylogenetic metrics that use branch length as a proxy for the evolutionary contribution of species are naturally sensitive to variations in tree topology (Calderón del Cid et al. 2024). In our case, such variations may arise both from the random resolution of soft polytomies and from pruning the trees to depict only the species present in local assemblages. Therefore, all metrics and analyses accounting for evolutionary history should be interpreted as local metrics (indicated herein with a subscript) rather than global values. We further discuss the limitations of this approach in section *4.5. Conservation from an evolutionary perspective*.

^2^In the Pacific, β values tend to be higher than those in other regions because of the occurrence of a single marine species (*Hydrophis platurus*). To enhance the pattern visualization in the figure, we plotted marine and continental communities separately and then piled them up. Therefore, sea and land max values are different.

