## Supplementary material for "Taxonomic, geographic, and phylogenetic patterns in the conservation status of the squamate reptiles (Reptilia: Squamata) of Colombia": Tables S3, S4, S5.docx

**Supplementary Files: Tables**

**Table S3.** Moran’s I coefficients (based on 999 permutations). Significant values are highlighted in bold.

| **Diversity** | **Moran’s I^*^** | ***p-value*** |
| --- | --- | --- |
| Species richness (α) | 0.91 | **0.001** |
| Species turnover (β) | 0.90 | **0.001** |
| Threatened species | 0.48 | **0.001** |
| Endemic species | 0.84 | **0.001** |
| Phylo. Endemism | 0.70 | **0.001** |

^*^Calculated using a queen’s scheme and the zero-policy set to TRUE for pixels without neighbors.

**Table S4.** Performance metrics for the relationship between elevation and squamate diversity in Fig. 3. The best models are highlighted in bold.

|  | **OLS** | | | | **Squared OLS** | | | |  |
| --- | --- | --- | --- | --- | --- | --- | --- | --- | --- |
|  | **R^2^** | **RMSE** | ***p-value*** | **AIC** | **R^2^** | **RMSE** | ***p-value*** | **AIC** | **ΔAIC** |
| Species richness (α) | 0.23 | 31.48 | < 0.001 | 75424 | **0.23** | **31.38** | **< 0.001** | **75380** | **1** |
| Species turnover (β) | 0.31 | 0.10 | < 0.001 | -14088 | **0.32** | **0.10** | **< 0.001** | **-14215** | 276 |
| Threatened species | 0.06 | 0.39 | < 0.001 | 7491 | **0.08** | **0.39** | **< 0.001** | **7319** | 254 |
| Endemic species | 0.09 | 3.67 | < 0.001 | 42136 | **0.13** | **3.58** | **< 0.001** | **41742** | 327 |
| Phylo. Endemism | 0.09 | 8.47 | < 0.001 | 55096 | **0.12** | **8.30** | **< 0.001** | **54772** | 258 |

**Table S5.** AIC values for the spatial autoregressive models fitted. The best models are highlighted in bold.

|  | **SAR_lag_** | | | **SAR_error_** | | |  |
| --- | --- | --- | --- | --- | --- | --- | --- |
|  | **R^2*^** | **RSME** | **AIC** | **R^2*^** | **RSME** | **AIC** | **ΔAIC** |
| Species richness (α) | **0.94** | **7.90** | **6514** | 0.94 | 7.43 | 6539 | 25 |
| Species turnover (β) | 0.84 | 0.06 | -2442 | **0.87** | **0.05** | **-2615** | 173 |
| Threatened species | 0.41 | 0.69 | 268 | **0.41** | **0.68** | **268** | 0 |
| Endemic species | 0.86 | 2.37 | 1613 | **0.86** | **2.27** | **1606** | 7 |
| Phylo. Endemism | 0.60 | 5.72 | 5819 | **0.65** | **4.93** | **5705** | 114 |

^*^Because spatial analyses do not provide standard R^2^ values, a Nagelkerke pseudo-R^2^ value is calculated.
